# Heart rate and its relationship with activity in free-ranging Cheloniidae sea turtles

**DOI:** 10.1101/557884

**Authors:** Junichi Okuyama, Maika Shiozawa, Daisuke Shiode

**Author notes:** Junichi Okuyama (Correspondence), Address: Research Center for Subtropical Fisheries, Seikai National Fisheries Research Institute, Japan Fisheries Research and Education Agency, Ishigaki, Okinawa, 907-0451 Japan.

## Abstract

The primary oxygen stores in Cheloniidae sea turtles are in the lungs. Therefore, management of blood oxygen transportation to peripheral tissues by cardiovascular adjustments while diving is crucial to maximize benefits from dives. However, heart rate, particularly cardiac response to exercise in free-ranging dives, has rarely been examined for sea turtles. In this study, heart rate and its relationship with the amount of activity were determined in six free-ranging green turtles using bio-logging techniques. Our results demonstrated that resting heart rate took 7–11 h to reduce to steady levels after turtles were released in the tank, indicating that turtles may not present normal physiological rates right after release. After heart rate reduction, resting heart rate of green turtles in free-ranging dives was generally low (10.9 ± 2.5 bpm), but they often presented arrhythmia (4–54 bpm) even in resting states. The amount of activity during a dive linearly increased heart rate, but maximum heart rates (39.0–69.8 bpm) were recorded during ventilation at surface. These results indicate that turtles have the capability of cardiac response to increased metabolic demands in their muscles while submerged, and also of cardiovascular adjustment for a rapid renewal of oxygen stores and removal of CO_2_ during ventilation. Such well-organized cardiac adjustments may be because of characteristics of Cheloniidae sea turtles such as ectothermy and oxygen storage in lungs while submerged.

**Summary statement:** Green sea turtles in free-ranging dive had generally lower heart rate compared to other air-breathing divers and it varied with the amount of exercise. Turtles often showed extreme arrhythmia.

## Introduction

Many air-breathing vertebrates are known to exhibit bradycardia when submerged, which allows them to regulate their blood oxygen depletion rate, thereby conserving onboard oxygen stores (Kooyman, 1989; Butler and Jones, 1997). This physiological mechanism enables these animals to submerge for prolonged periods (Kooyman, 1989; Butler and Jones, 1997). However, a paradoxical situation is created because underwater activity, such as swimming, should promote an elevation in heart rate to support the increased demands of the aerobic metabolism if metabolism is to remain aerobic. Thus, the potential for conflict between diving bradycardia and exercise responses has been investigated for marine mammals (Davis and Williams, 2012; Noren et al., 2012; Williams et al., 2015). These studies have revealed that marine mammals modify their level of bradycardia according to the amount of activity while submerged (Davis and Williams, 2012; Noren et al., 2012; Williams et al., 2015). Moreover, a recent study revealed that bradycardia was altered not only by exercise, but also by dive depth and the possibility of lung collapse (Williams et al., 2015).

Sea turtles are ectothermic marine animals and have well-adjusted physiological functions for prolonged dives (reviewed by Lutcavage and Lutz, 1997; Williard, 2013). Their considerably slow metabolism, for instance, enables turtles to undergo extremely prolonged dives (e.g., Hochscheid et al., 2005). Moreover, in sea turtles, particularly in Cheloniidae species, the primary oxygen stores are in the lungs (> 70% of the oxygen present in whole body) rather than in the blood or tissues (Lutz and Bentley, 1985; Lutcavage and Lutz, 1997). Therefore, the management of blood oxygen transportation to peripheral tissues (e.g., muscle) while diving by cardiovascular adjustments is crucial to maximize dive duration and the benefits of underwater activities, such as feeding, predator avoidance, and mating. Therefore, the cardiac responses of sea turtles to diving have been of particular interest to comparative biologists (Berkson, 1966; Davenport et al., 1982; Butler et al., 1984; West et al., 1992; Southwood et al., 1999). However, all of these previous studies, except Southwood et al. (1999), were conducted under experimental conditions in a limited space, and heart rate in free-ranging sea turtles has rarely been examined. Moreover, unlike for marine mammals, knowledge on sea turtle bradycardia while diving remains limited. In this context, the objectives of this study were to examine general aspects of heart rate in free-ranging green sea turtles, *Chelonia mydas* (Linnaeus 1758) and to determine their cardiac response to activity while diving.

## Materials and methods

### Experimental protocol and instruments

Our experiments were conducted in a tank (H × L × W = 10 m × 10 m × 2.2 m) at the Research Center for Subtropical Fisheries, Seikai National Fisheries Research Institute, Japan Fisheries Research and Education Agency, Japan. Six wild green turtles were captured under the permission of Okinawa Prefecture (No. K30-2) (Table 1).

**Table 1.** Summary of physiological characteristics of turtles and experimental conditions.

| ID | SCL (cm) | BW (kg) | Duration of<br>experiment (h) | Duration before resting<br>HR<sub>min</sub> reduction (h) | T<sub>w</sub> (°C) |
| --- | --- | --- | --- | --- | --- |
| CM1 | 66.3 | 34.5 | 16.3 | 8 | 29.5 - 30.3 |
| CM2 | 58.1 | 22.5 | 19.5 | 10 | 28.9 - 29.8 |
| CM3 | 58.1 | 22.4 | 19.1 | 9 | 29.1 - 29.9 |
| CM4 | 49.5 | 15.2 | 20.2 | 11 | 28.3 - 29.4 |
| CM5 | 48.7 | 13.2 | 21.5 | 10 | 29.2 - 29.9 |
| CM7 | 42.3 | 8.5 | 17.2 | 7 | 28.7 - 29.5 |

Electrocardiogram (ECG) data of free-ranging green turtles were recorded using an ECG data logger (W400-ECG, Little Leonard Co., Tokyo, Japan, 21 mm × 109 mm cylindrical logger, 5 ms sampling interval, voltage range ± 5.9 mV, 60 g, 2 GB memory). Three wires extending from the ECG logger were soldered to two stainless-steel electrodes (positive and negative, 0.9 mm in diameter, 30–35 mm in length) and a 21–23 gauge sterile needle that functions as a ground connection. Length of the two electrodes was modified according to the size of turtles to record clear heart beats. These electrodes were placed in the plastrons under anesthesia. Medetomidine (0.2 ml/kg of body weight, Fujita Pharmaceutical Co., Ltd., Tokyo, Japan) was used as a sedative and injected into the base of front flippers. Approximately five minutes after sedative injection, anesthesia was initiated by slow injection of 1-ml diluted water with Propofol (0.01 ml/kg; Mylan Inc., PA, USA) into the cervical vein. Two ECG electrodes were placed through a 1.5-mm hole in the plastron over the heart (Fig. 1). Dental acrylic (Fuji IX GP; GC Co., Ltd., Tokyo, Japan) was used to anchor the electrode and seal the hole. A ground electrode was inserted immediately anterior to the hindlimb at a depth of 55 mm. Electrode placement required less than 15 min. The ECG data logger was placed on the carapace using epoxy putty (Cemedine Co., Ltd., Tokyo, Japan). Then, three wires were fixed on the carapace and plastron and covered using a silylated urethane resin (Konishi Co., Ltd., Osaka, Japan) and duct tape (Koyo Kagaku Co., Ltd., Tokyo, Japan) to prevent electrical noise. After attachment procedures for electrodes and loggers, atipamezole (0.3 ml/kg Mepatia; Fujita Pharmaceutical Co., Ltd.) was injected into the base of front flippers as a medetomidine antagonist, to allow the turtle to recover from sedation. Then, each turtle was maintained in the plastic container (H × L × W = 1 m × 0.7 m × 0.2 m) for ∼1–1.5 h before the epoxy putty and resins completely hardened.

**Fig. 1.**
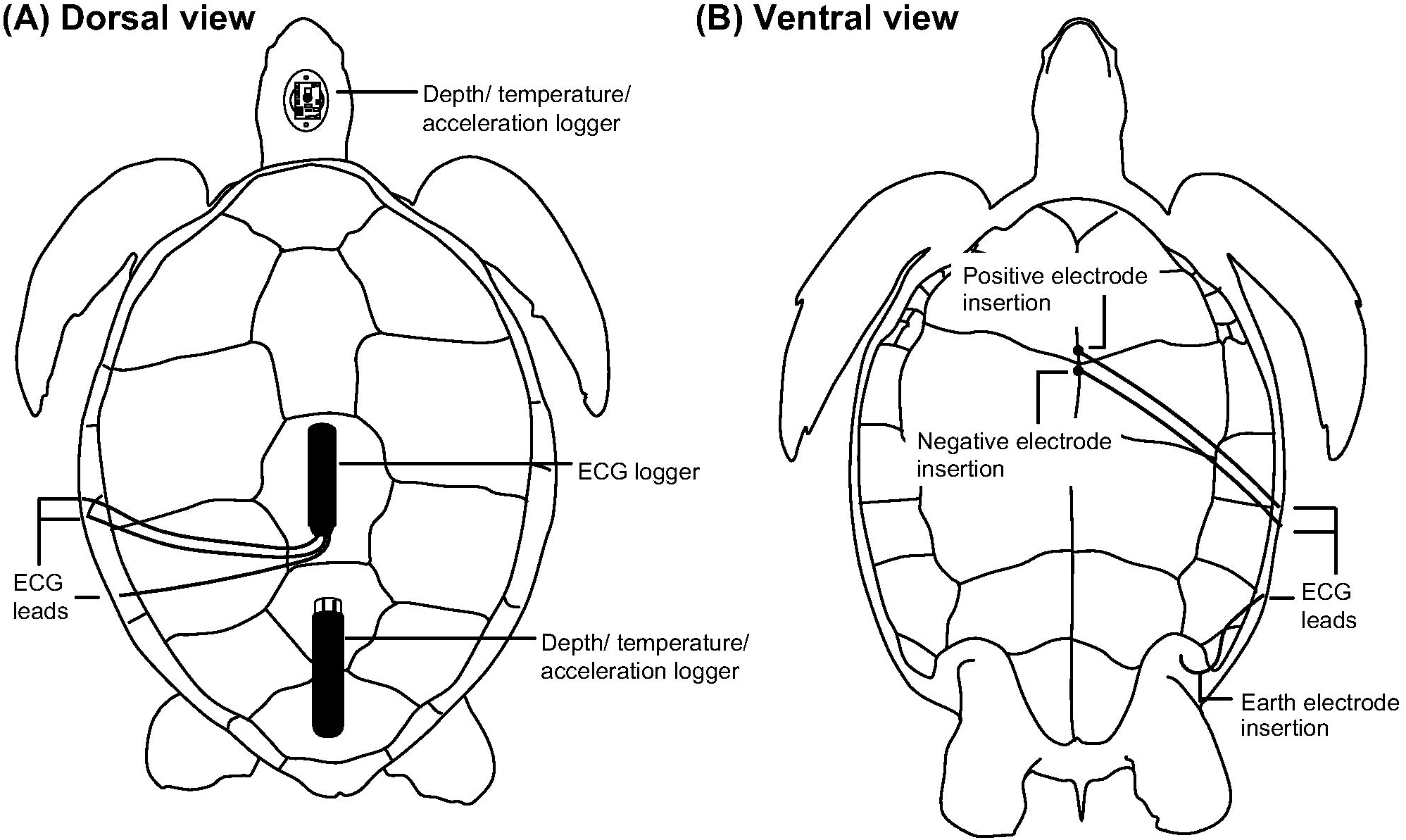
Illustrations of data logger attachments to a green turtle. (A) Dorsal view showing the placement of the data loggers and of the electrode leads on the carapace. (B) Ventral view showing the placement of three electrodes and leads on the plastron.

In addition to heart rate, depth, temperature, and 3-axis acceleration were simultaneously recorded using a multi-channel data logger (ORI400-D3GT: 12 mm diameter, 45 mm length, 9 g in air, memory capacity of 11.5 million data points; or W380-PD3GT: 21 mm diameter, 114 mm length, 59 g in air, memory capacity of 16 million data points; Little Leonardo Co.) placed on the carapace using epoxy putty (Fig. 1). It recorded depth and temperature at 1 s intervals and 3-axis accelerations at 1/20 or 1/16 s intervals. The maximum range of the depth sensor was 400 or 380 m, with a resolution of 0.1 m. The measurement range of the accelerometers was ± 3 or 4 *g*, with a resolution of 0.00009 or 0.002 *g*. Acceleration data were separated into two components: the high frequency component, which was a dynamic acceleration representing turtle movement (such as swimming), and the low frequency component, which was gravity acceleration representing the turtle’s body angle (Okuyama et al., 2012). This frequency separation was conducted by filtering analysis (see Okuyama et al. [2012] for details). Moreover, acceleration data were used to calculate the amount of activity performed by each turtle, which was given by overall dynamic body acceleration (ODBA; Wilson et al., 2006). ODBA was used as a proxy of the amount of activity and, consequently, metabolic rate in sea turtles (Halsey et al., 2011; Okuyama et al., 2014). ODBA was calculated from the high frequency component of the 3-axis acceleration data. Furthermore, the number of breaths at the surface after each dive was counted by the head-mounted acceleration data logger (G6a+; dimensions: 40 mm × 28 mm × 17 mm, weight: 19 g in air; measurement range: ± 2 *g* with a resolution of 0.002 *g*, Cefas Technology Limited, Suffolk, UK; Fig. 1). Head angle of sea turtles was calculated by the gravity component of acceleration data and depth data and allowed us to count the breathing behavior (Okuyama et al., 2010). Briefly, breathing behavior was defined as occurring when turtles raised their head up with angle > 30° at a depth < 0.15 m (Okuyama et al., 2010).

Each turtle with the dataloggers was released into the tank and allowed to swim freely for 16.3–21.5 h. Temperature during the experiments ranged from 28.3–30.3°C (Table 1). After the experiments, the turtles were caught and immediately removed from the tank. Then, blood samples were collected from the cervical vein using a syringe (1 ml, Nipro Co., Osaka, Japan) and a needle (20-gage, Terumo Co., Tokyo, Japan). Partial pressures of oxygen and CO_2_, besides lactate level in the venous blood, were measured using an i-STAT 1 Analyzer (Abbott Point of Care Inc., IL, USA) to be used as indicators of whether the turtles engaged in anaerobic dives during the experiments. After blood sampling, all electrodes were removed, and the holes were filled with dental acrylic. The turtles were maintained in the rearing tank while the recovery condition of the holes was observed. Two weeks after the experiments, the holes were completely closed, and the turtles were released back into the sea.

### Extraction of heart rate from ECG data and data analysis

Data of ECG, depth, and acceleration were analyzed using Igor Pro version 6.36 (Wavemetrics, OR, USA). As ECG waveforms and the amplitude of QRS peaks changed in each situation (see the Results section), heart rate was used only to investigate the turtles’ cardiac response. The R peak in the QRS complex (the main graphical deflections in an ECG tracing; Fig. 2) was easily detectable, and thus was regarded as one heartbeat.

**Fig. 2.**
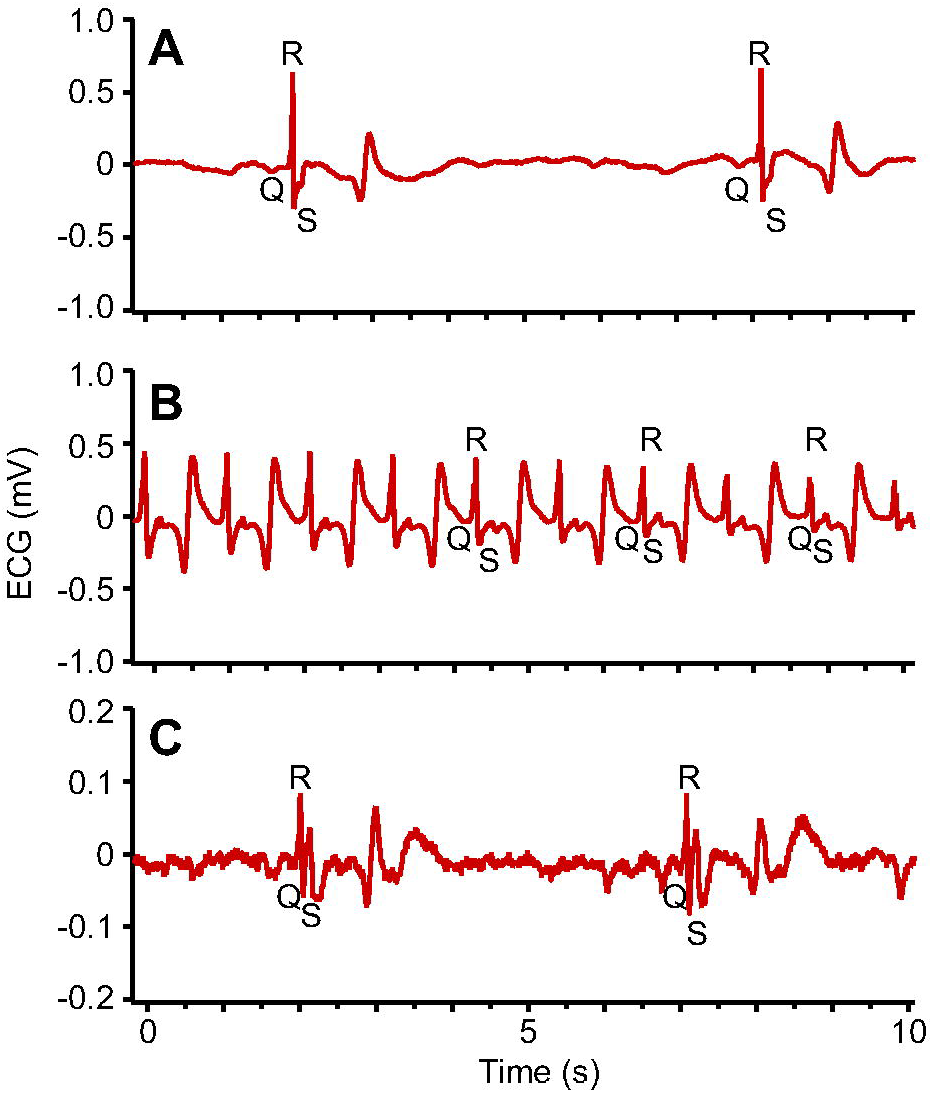
Typical examples of electrocardiogram (ECG) traces showing heart beats in green turtles. (A) Normal ECG trace showing clear a QPR complex. (B) ECG trace at extreme tachycardia. (C) ECG trace when recording condition was not well. Note that the range of the vertical axis is different from those of the other two.

Heart rate was analyzed in two time-scales. One was HR_*dive*_, which was calculated by the number of heartbeats during a dive divided by dive duration (min) to investigate the relationship of heart rate between dives and post-dive surface duration. A dive was defined as the period when turtles submerged deeper than 0.5 m and for longer than 30 s; the remaining period was regarded as surface period. The other time-scale was HR_*min*_, which was calculated by the number of heartbeats per minute for the analysis of detailed cardiac response. However, when HRs_*min*_ included the surface, pre-surface ascend, and post-surface descend phases, such data were excluded from the analysis to eliminate the effect of tachycardia at surface. Moreover, in some cases, the QRS complex was not identified because of the noise (see the Results section). Therefore, the data of HRs_*min*_ and HRs_*dive*_ including such noise were excluded from the analysis.

To investigate resting heart rate, and also to understand the relationship between heart rate and activity level, behavior was divided into two states every minute: resting state and active state. Resting state was defined as the behavior when dynamic acceleration, body angle, and depth were constant throughout a minute, representing a turtle stayed still at the bottom of tank. All other periods were considered to be an active state. Similarly, resting dive was defined as dives in which the turtles were almost in the resting states at the bottom phase of dives. The remaining dives were defined as active dives.

### Statistical analysis

To determine the baseline of heart rate in green turtles, we analyzed the transition of resting HRs_*min*_, because these values appeared to decrease with time elapsed after the release (see the Results section). Thus, HRs_*min*_ data were summarized every hour after the release. Statistical outliers in each hour-dataset were excluded using the Smirnov-Grubbs test. The remaining non-outlier data were analyzed to determine the significant difference in HRs_*min*_ among hours after the release using multiple comparison Tukey-Kramer test, based on the hypothesis that the baseline of resting heart rate is maintained at a constant level with no significant difference among hours when the heart rate is reduced.

A generalized linear mixed model (GLMM) with a Poisson distribution and a log link function was used to determine the factors affecting heart rate (HR_*min*_). Mean ODBA during a minute, body weight of turtles, and mean water temperature were treated as explanatory variables. Individual was treated as a random effect. To determine the effect of heart rate on dive duration, we also investigated the factors regulating dive duration using a GLMM with a Gamma distribution and a log link function. For this model, heart rate (HR_*dive*_), mean ODBA, mean water temperature during each dive, and body weight were treated as explanatory variables. We used the ‘lme4’ package in the R v. 3.5.2 (R Development Core Team, 2018) software to run the GLMM analyses.

## Results

### Extraction of heart rate

A total of 113.8 h of depth, temperature, 3-axis acceleration, and ECG data were obtained from six turtles (Table 1). Most of the ECG data showed a clear QRS complex, but waveform and amplitude of the QRS peaks changed for each situation, presumably because the relative positions of the two electrodes toward the heart slightly changed as a result of the active movements of the turtles (e.g., swimming; Fig. 2). Therefore, in this study, only heart rate was used for the analysis. Furthermore, in some cases, the QRS complex could not be distinguished because of the noise; these data were excluded from the analysis.

Turtles often rested at the bottom of tank, and they swam around the tank during the remaining period. Immediately after turtle release, the resting HRs_*min*_ were higher than those in other periods, and then they gradually decreased (Fig. 3). Multiple comparison tests revealed that the resting HRs_*min*_ in each hourly data did not significantly differ between all comparisons from 7–11 h after the release of each turtle (Fig. 3, Table 1). The statistical results of the multiple comparison tests are shown in Table S1. We therefore regarded periods with non-significant difference in HRs_*min*_ as the condition in which the heart rates were reducing. To correctly determine the heart rates of sea turtles under natural conditions, hereafter, only the data from after heart rate reduction were used in the analyses. Thus, 1,875 minute-datasets including HRs_*min*_, mean ODBA, and mean water temperature from six turtles were used in this study (Table 2). Moreover, 347 dive-datasets including HRs_*dive*_, dive duration, mean ODBA, and mean water temperature, as well as post-dive surface data (HRs_*surface*_, the number of breaths) were also used (Table 3).

**Table 2.**
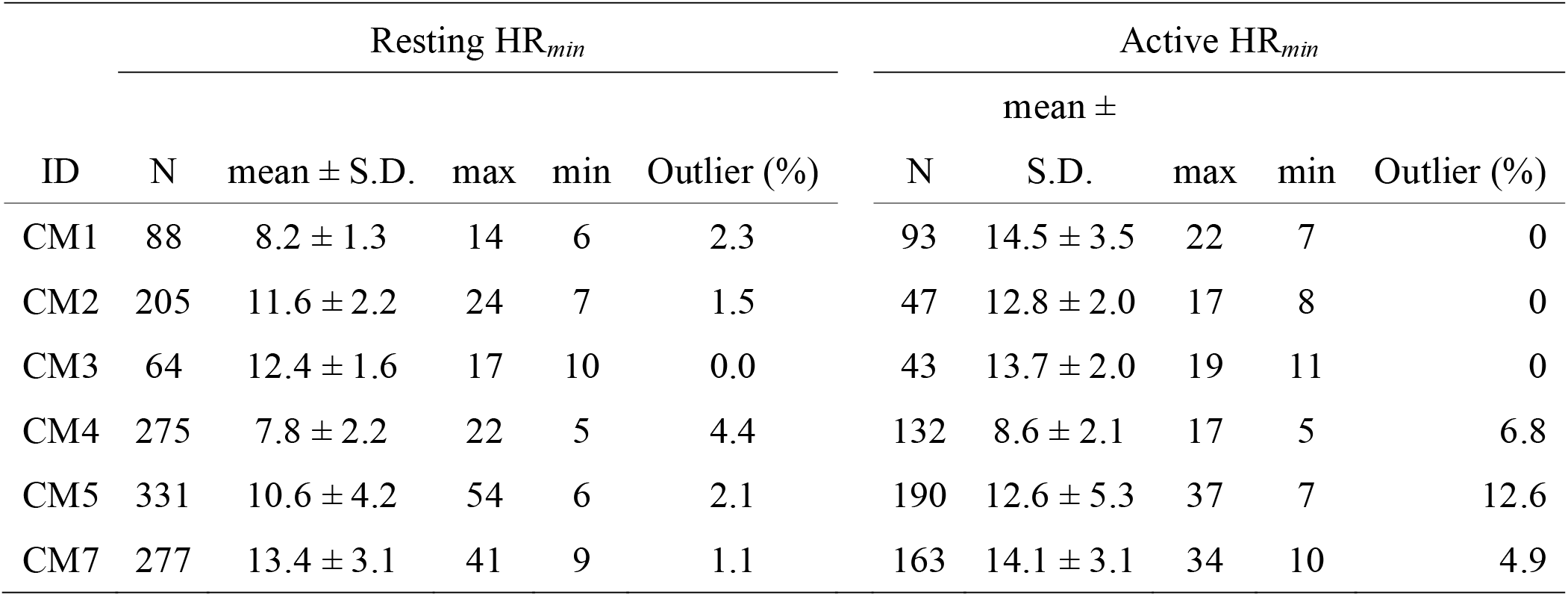
Summary of minute-heart rate in each turtle.

**Table 3.** Summary of dive-heart rate in each turtle.

| Resting dive |  |  |  |  | Active dives |  |  |  |
| --- | --- | --- | --- | --- | --- | --- | --- | --- |
| ID | N | Dive | Post-dive at surface |  | N | Dive | Post-dive at surface |  |
|  |  | HR<sub>dive</sub><br>(bpm) | HR<sub>surface</sub><br>(bpm) | N.<br>breaths |  | HR<sub>dive</sub><br>(bpm) | HR<sub>surface</sub><br>(bpm) | N.<br>breaths |
|  |  | (max - min) | (max - min) | (max - min) |  | (max - min) | (max - min) | (max - min) |
| CM1 | 10 | 9.4 ± 0.7 | 38.2 ± 7.0 | 1.6 ± 1.2 | 95 | 17.7 ± 3.3 | 28.4 ± 5.1 | 1.0 ± 0.2 |
|  |  | (10.2 - 8.2) | (52.0 - 30.0) | (5 - 1) |  | (30.4 - 10.7) | (42.0 - 12.0) | (2 - 1) |
| CM<br>2 | 7 | 12.9 ± 2.1<br>(17.0 - 11.3) | 51.1 ± 9.2<br>(69.8 - 30.7) | 1.1 ± 0.4<br>(3 - 1) | 7 | 23.4 ± 5.7<br>(31.7 - 16.5) | 45.5 ± 8.1<br>(60.6 - 18.4) | 1.0 ± 0.1<br>(2 - 1) |
| CM<br>3 | 2 | 14.6 ± 0.5<br>(14.9 - 14.3) | 32.2 ± 4.1<br>(39.0 - 27.7) | 1.0 ± 0.2<br>(2 - 1) | 4 | 15.4 ± 1.1<br>(16.9 - 14.3) | 35.0 ± 5.7<br>(44.1 - 28.8) | 1.0 ± 0.1<br>(2 - 1) |
| CM<br>4 | 2<br>0 | 8.0 ± 0.5<br>(9.2 - 7.3) | 28.0 ± 8.1<br>(49.8 - 17.8) | 1.4 ± 0.9<br>(4 - 1) | 2<br>5 | 11.1 ± 5.0<br>(32.5 - 7.5) | 23.0 ± 5.5<br>(39.9 - 17.3) | 1.0 ± 0.1<br>(2 - 1) |
| CM<br>5 | 3<br>1 | 11.3 ± 1.7<br>(18.3 - 9.9) | 32.9 ± 8.2<br>(58.8 - 19.6) | 1.4 ± 0.7<br>(3 - 1) | 5<br>3 | 15.7 ± 4.9<br>(30.2 - 9.7) | 26.5 ± 7.2<br>(49.0 - 15.7) | 1.0 ± 0.2<br>(2 - 1) |
| CM<br>7 | 3<br>4 | 15.0 ± 2.0<br>(22.6 - 12.9) | 47.8 ± 8.4<br>(69.0 - 35.2) | 2.5 ± 1.4<br>(6 - 1) | 5<br>9 | 22.3 ± 8.6<br>(45.2 - 13.8) | 38.0 ± 5.2<br>(50.6 - 27.7) | 1.2 ± 0.5<br>(3 - 1) |

**Fig. 3.**
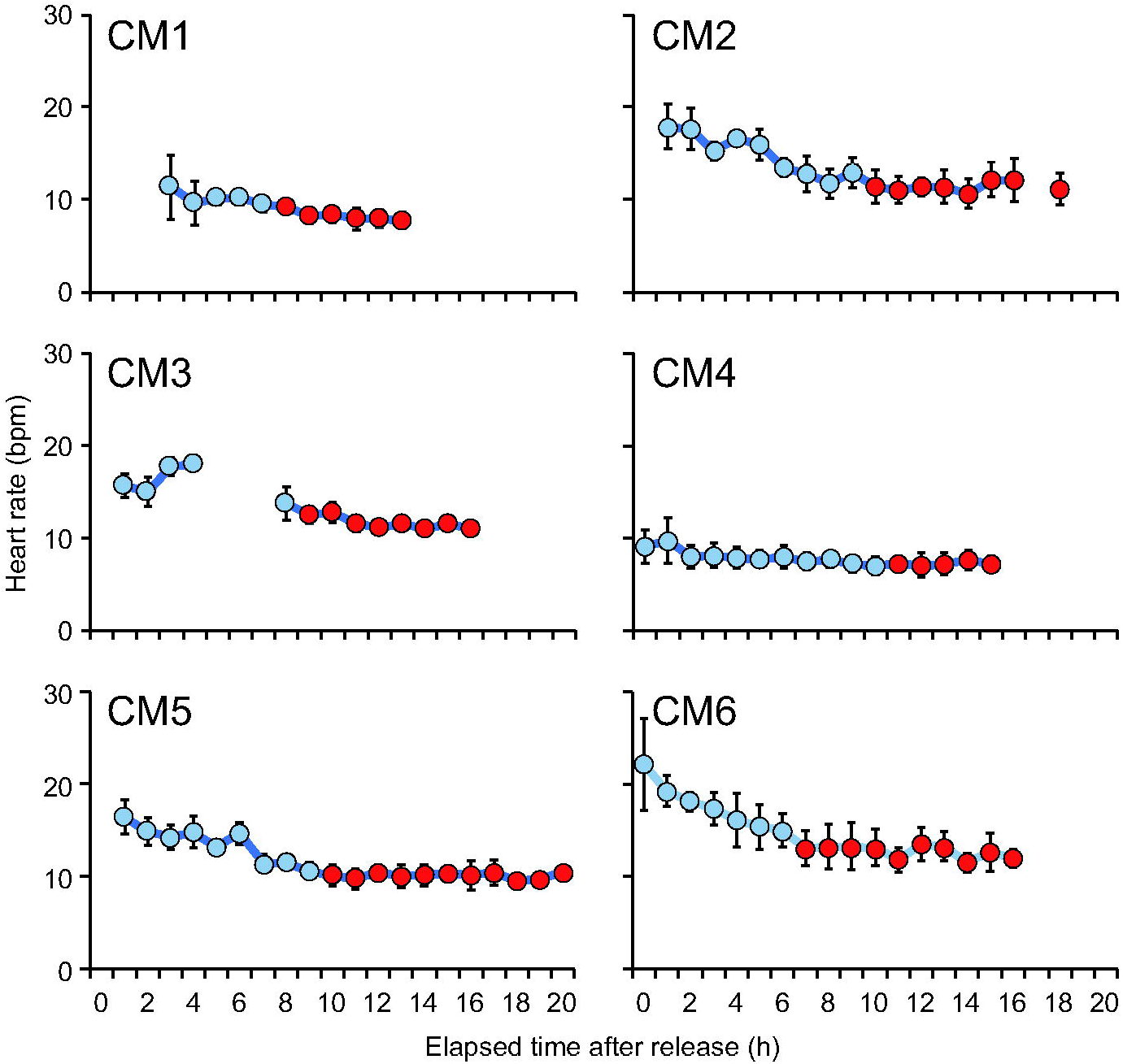
Heart rates in the resting states with elapsed time after release into the tank in each turtle. Circles and vertical bars represent the mean value and standard deviation of heat rate in the resting state in each hour, respectively. Statistical outliers were excluded. Red colors represent heart rate reduction identified using a multiple comparison test.

### Heart rate

Resting HR_*min*_ was 10.9 ± 2.5 bpm on average (± s.d.) (Table 2). However, it was noteworthy that resting HRs_*min*_ was not constant even after it reduced; abnormal events of instant, extreme tachycardia (54 bpm at maximum) sometimes occurred and were detected by the statistical outlier test (0.0–4.4% of the resting HRs_*min*_; Table 2, Fig. 4).

**Fig. 4.**
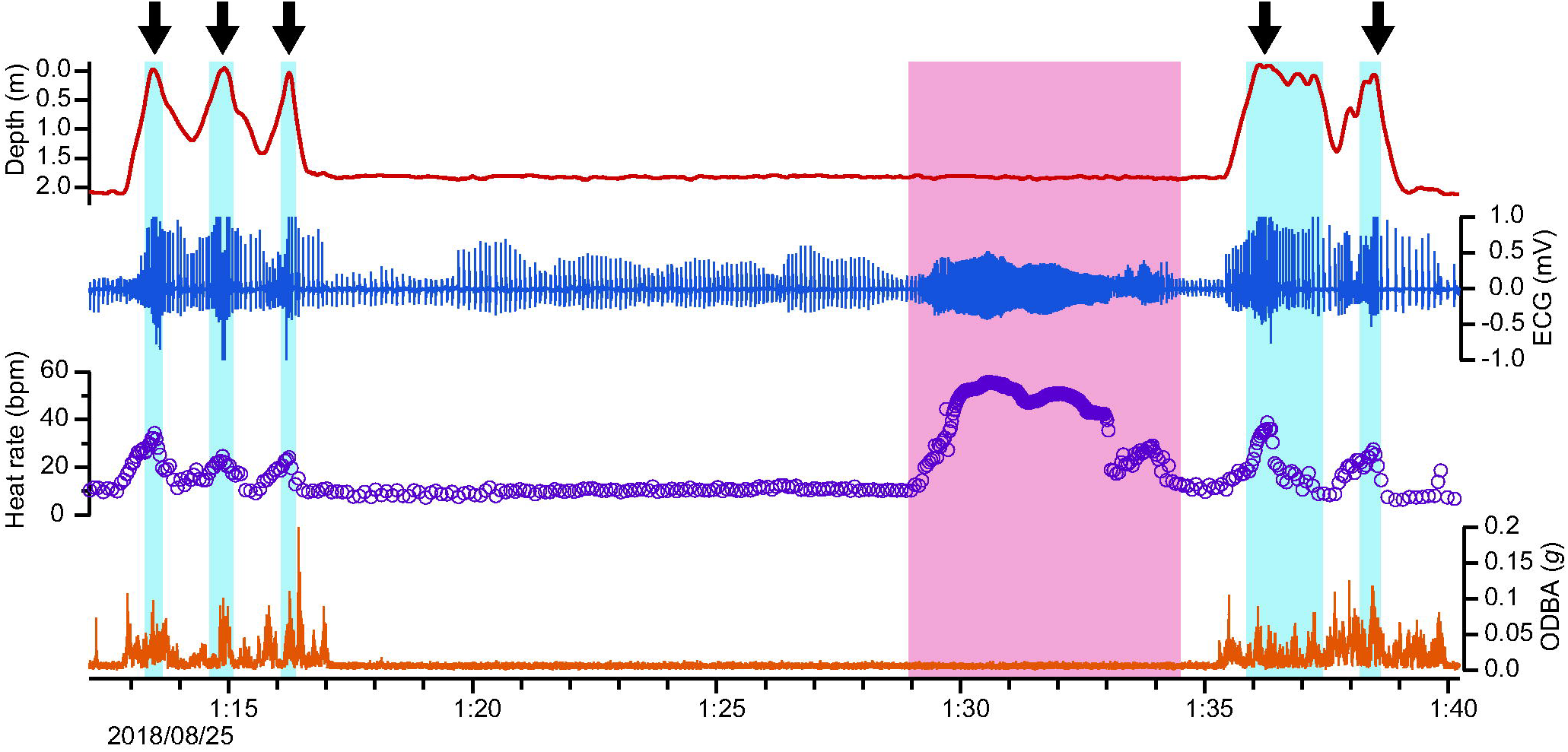
Typical cardiac responses during a resting dive in a green turtle (CM5). Blue and red backgrounds represent surface periods and extreme tachycardia period. Black arrows represent breathing event.

Active HRs_*min*_ was 12.7 ± 2.1 bpm on average (± s.d.) (Table 2). The GLMM analysis showed the HRs_*min*_ significantly increased with mean ODBA (Fig. 5), but had no significant relationship with body size and water temperature (Table 4).

**Table 4.** Summary of the statistics from two generalized linear mixed models’ analyses.

| Response variable | Distribution | Explanatory variables | Estimate | Std. Error | *z or t* | *P* |
| --- | --- | --- | --- | --- | --- | --- |
| Heart rate | Poisson | mean ODBA | 0.10 | 0.01 | 13.06 | <0.001 |
|  |  | Body weight | -0.019 | 0.069 | -0.27 | 0.785 |
|  |  | Temperature | -0.01 | 0.02 | -0.69 | 0.493 |
|  |  | Intercept | 2.43 | 0.08 | 30.22 | <0.001 |
| Dive duration | Gamma | Heart rate | -0.53 | 0.04 | -14.33 | <0.001 |
|  |  | Mean ODBA | -0.28 | 0.04 | -8.00 | <0.001 |
|  |  | Body weight | -0.20 | 0.09 | -2.20 | <0.05 |
|  |  | Temperature | 0.01 | 0.09 | 0.07 | 0.945 |
|  |  | Intercept | 1.25 | 0.05 | 23.43 | <0.001 |

**Fig. 5.**
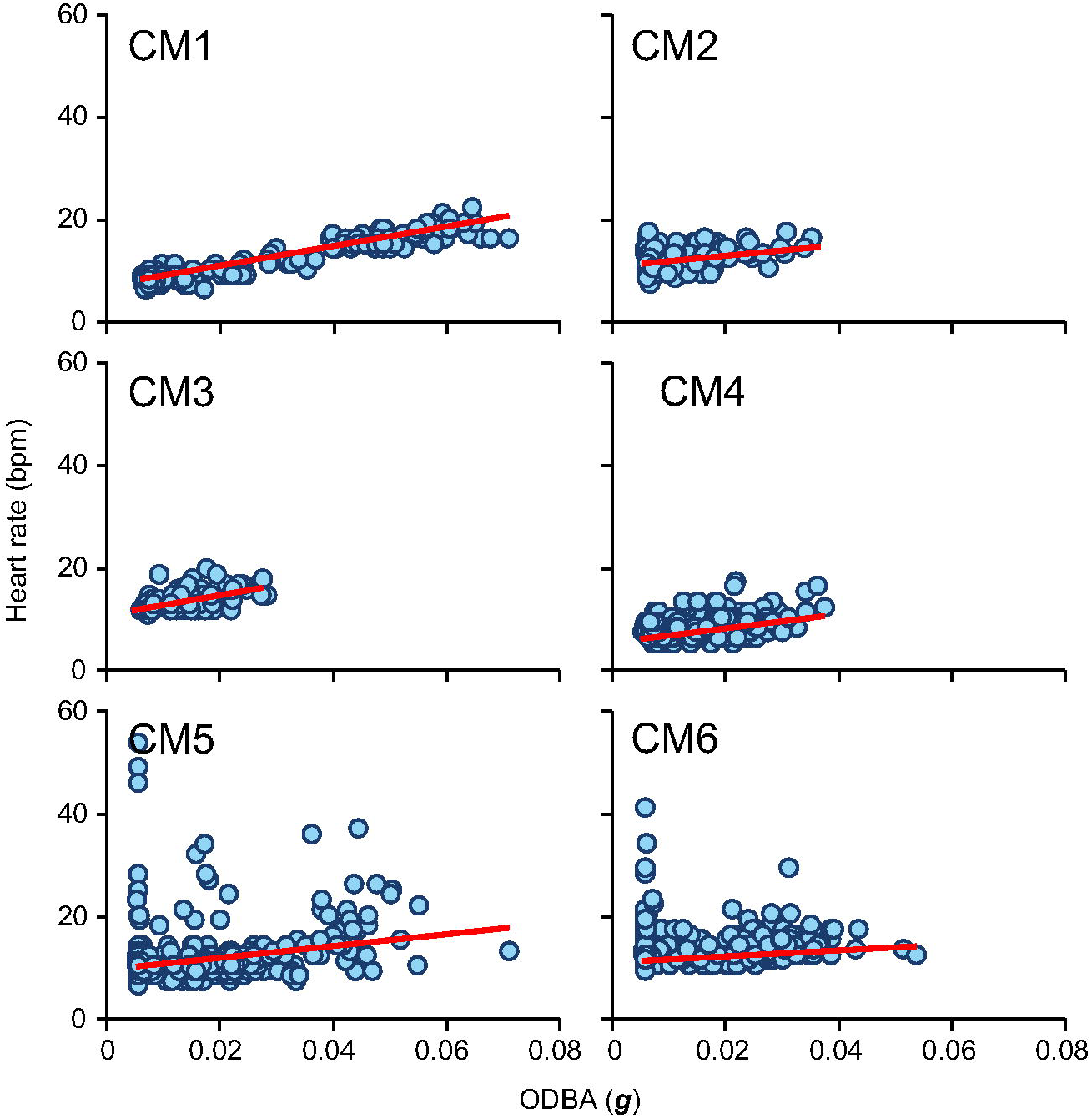
Relationships between heart rate and the overall dynamic body acceleration (ODBA) in each turtle after heart rate reduction. Red lines represent the linear regression line calculated using a least-square method.

### Cardiac response to the diving behavior

Dive duration was 10.2 ± 2.2 min and 2.9 ± 1.0 min for resting and active dives, respectively. Clear bradycardia was observed in all dives. Heart rate began to decrease immediately after the start of submergence, and it increased concurrently with the start of ascending to surface (Fig. 4). Thus, HRs_*dive*_ was always higher than HRs_*min*_, because it included such ascending and descending periods in which heart rates elevated because of tachycardia during surface period. Resting HRs_*dive*_ was 11.9 ± 2.8 bpm (Table 3). Mean resting HRs_*surface*_ was 38.4 ± 9.2 bpm and increased 2.2–4.1-fold from resting HRs_*dive*_ (Table 3). Meanwhile, mean active HRs_*dive*_ was 17.6 ± 4.6 bpm, and mean active HRs_*surface*_ was 32.7 ± 8.3 bpm (Table 3). Resting HRs_*surface*_ was significantly higher than active HRs_*surface*_ (paired *t*-test, N = 6, *t*_5_ = 3.0, *P* < 0.05). Number of breaths during post-dive surface duration was 1.5 ± 0.5 in resting dives and 1.0 ± 0.1 in active dives (Table 3).

### Effect of heart rate on dive duration

The GLMM analyses for all dive data, including those on before heart rate reduction, revealed that dive duration was significantly affected by heart rate, mean ODBA, and body weight of turtles, but not by water temperature (Table 4). The average (± s.d.) partial pressures of oxygen and CO_2_ and lactate level in the venous blood immediately after the experiments were 51.3 ± 9.2 mmHg, 66.3 ± 9.4 mmHg, and 1.0 ± 1.2 mmol L^-1^, respectively.

## Discussion

This study described the heart rate of green turtles in natural conditions, and the changes in this feature caused by various factors, such as amount of activity, and air ventilation at surface. Sea turtles are regarded as ‘surfacers’ rather than ‘divers,’ because they spend most of their time underwater and surface only briefly for gas exchange (Kooyman, 1989). Therefore, bradycardia is a usual condition in the life history of sea turtles. The assessment of heart rate under normal condition in voluntary dives, as performed in this study, is an essential approach to understand the diving physiology of sea turtles. Our study demonstrated that the heart rate of green turtles was considerably lower than those of leatherback turtles (Southwood et al., 1999), and marine mammals and birds (Butler and Jones, 1997). On the other hand, the extent of tachycardia at surface in green turtles was wider than that in other animals (e.g., leatherback turtles, 1.4-fold [Southwood et al., 1999]; northern elephant seals *Mirounga angustirostris*, 1.56-fold [Andrews et al., 1997]; and bottlenose dolphin *Tursiops truncates*, 2.63-fold [Noren et al., 2012]). These well-organized cardiac adjustments may be a result of characteristics of Cheloniidae sea turtles such as being ectotherms with lungs with oxygen stores while submerged. Furthermore, these findings provide essential knowledge for understanding the green turtles’ diving strategy as well as their diving physiology.

### General perspective of heart rate in green turtles

Heart rate reduction after the animals were released in the tank took 7–11 h. This indicates that the turtles may have taken some time to reach their normal physiologically rates after release, even though they were observed to have settled on the bottom of the tank. Heart rate is known to increase in response to stress in turtles (Cabanac and Bernieri, 2000) and in many other taxa (Tarlow and Blumstein, 2007). Therefore, the stress caused by experimental handing including electrode attachment (although the turtles were under anesthesia) and by being released into a new environment (experimental tank) may have increased heart rate. It took ∼7–11 h for the turtle to recover from this stress. This information should be taken into account by researchers when estimating metabolic rates of activity levels in sea turtles (e.g., Halsey et al., 2011) as the metabolic rate may be overestimated because of heart rate elevation resulting from a stress condition. In this study, duration of resting dives before heart rate reduction was shorter than that after heart rate reduction, which demonstrates that high heart rates make dive duration shorter even in resting states without any active metabolism.

Voluntary dives in sea turtles are generally completed with aerobic metabolism (Butler et al., 1984; Okuyama et al., 2014). The low concentrations of lactate acid in venous blood collected immediately after the experiments supported that the turtles mostly performed aerobic dives in our study (Lutz and Bentley, 1985). Our results demonstrated that heart rate in sea turtles increased linearly with the amount of activity even when submerged with bradycardia. This indicates that sea turtles increase oxygen delivery to active muscles by blood flow during a dive in order to meet metabolic demands, as in marine mammals (Fedak et al., 1988; Williams et al., 1999; Davis and Williams, 2012; Williams et al., 2015). The extent of tachycardia in active states was not higher than that during surface ventilation and during extreme tachycardia in resting state. This indicates that turtles may maintain oxygen delivery to some extent during active swimming. This capability of cardiac adjustment may enable turtles to maintain aerobic metabolism even during active dives.

Previous laboratory studies reported that the mean resting heart rates for juvenile green turtles was around 24 bpm (Butler et al., 1984; West et al., 1992). In terms of scaling, no significant effect of body size was detected in this study, probably because of the small sample size. However, including the data obtained by previous studies (Butler et al., 1984: N = 5, body weight = 1.2 kg, water temperature = 28.8°C; West et al., 1992: N = 10, body weight = 1.2 kg, water temperature = 28.0°C), the resting heart rate (*f*_resting_) decreased with larger body weight (BW), which was expressed by the following power equation using the least square regression (*r*^2^ = 0.85, one-way *ANOVA, F*_*1.7*_ = 81.0, *P* < 0.001. Fig. 6):

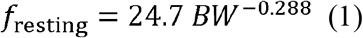

**Fig. 6.**
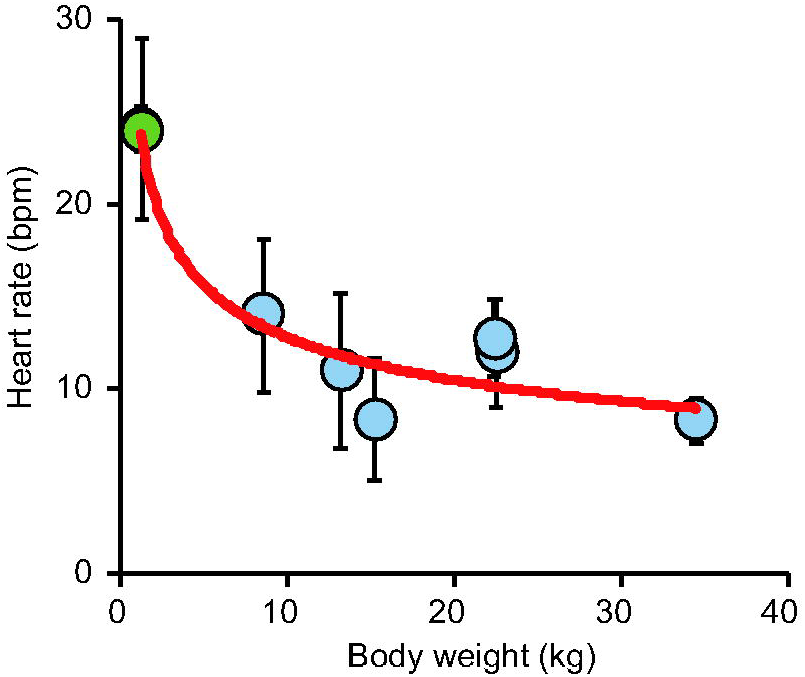
Heart rate in resting states (bpm) versus body weight (kg) in green turtles. Circles and verticals bars represent the mean values and standard deviations of heart rates. Blue circles represent data obtained in this study, whereas green circles represent data from Butler et al. (1984) and West et al. (1992). Red line represents the heart rate estimated from body weight using Eqn 1 (Heat rate = 24.7 BW^-0.288^).

Similar physiological parameters such as metabolic rate (Wallace and Jones, 2008) and aerobic dive limit (Hochscheid et al., 2007) were also reported to have a relationship with BW in sea turtles. Mass exponent (−0.288) of power regression of resting heart rate in green turtles are surprisingly similar to those in mammals (−0.25; Stahl, 1967) and in birds (−0.23; Calder, 1968). However, heart rate measured in shallow water chambers by previous studies (35–40 cm in depth; Butler et al., 1984; West et al., 1992) may be a bit higher than the results of our study, conducted in a large tank 2.2 m deep, because dive depth-induced lung collapse reduces heart rate in marine mammals (Williams et al., 2015).

Another factor affecting heart rate is ambient water temperature (Davenport et al., 1982); however, no significant effect was detected in this study, presumably because there were no substantial fluctuations in water temperature during the experiments (Table 1). Heart rates of free-ranging leatherback turtles during inter-nesting period in Costa Rica were documented as 17.4 bpm on average (Southwood et al., 1999). In that study, leatherbacks mostly conducted V-shaped dives. Such situation is likely similar to the active dives of the current study. Considering the large size difference between the adult leatherback turtles (250–400 kg; Southwood et al., 1999) and sub-adult green turtles (8.5–34.5 kg) in current study, heart rate in leatherback seems to be higher than that in green turtles. This might be because the body temperature of leatherbacks was somewhat higher than the ambient water temperature due to gigantothermy (Paladino et al., 1990).

### Abnormal cardiac rhythm in resting states

Our study found that abnormal cardiac rhythms including extreme tachycardia generally occurred in green turtles. Irregular cardiac rhythms were also reported in leatherbacks and green turtles (Davenport, 1982, Southwood et al., 1999) and in marine mammals (Williams et al., 2015). In the case of the marine mammals, the cardiac arrhythmias may have occurred as a result of the interplay between sympathetic and parasympathetic drivers for exercise and diving (Williams et al., 2015). This aspect may also be true for sea turtles. However, extreme tachycardia in resting states may also indicate that turtles may adjust the oxygen stored in peripheral tissues (e.g., locomotory muscles), which suffered from anoxia by temporarily increasing blood flow during a dive. Otherwise, it may represent a ‘fight-or-flight’ cardiac response toward some dangerous situation felt by the turtles (i.e., stress).

### Cardiac response to diving

Many diving mammals and birds increase their heart rate towards the end of a dive in anticipation of surfacing (Butler and Jones, 1997), because an anticipatory tachycardia might restore blood flow to peripheral tissues, flushing out metabolites that may have built up and allowing for a more efficient removal of metabolic by-products and uptake of oxygen during recovery time at the surface (Thompson and Fedak, 1993). Our study demonstrated that green turtles also rapidly elevated their heart rate at the begging of ascending to surface, as well as leatherback turtles (Southwood et al., 1999). Tachycardia observed during post-dive surface durations was quite faster than that observed for leatherback turtles (24.9 bpm, Southwood et al., 1999). Cheloniidae species use their lungs as the primary oxygen store while diving and have relatively larger lung tidal volumes than those of deep-dive leatherbacks turtle (Lutcavage and Lutz, 1991). These facts may allow a more rapid renewal of oxygen stores and removal of CO_2_ in Cheloniidae species.

Tachycardia during post-dive surface durations in resting dives was faster than in active dives. Okuyama et al. (2014) suggested that green turtles almost deplete their oxygen stored during resting dives, and then replenish their oxygen content at surface by taking several breaths (Table 3). Thus, faster tachycardia may indicate that green turtles make additional adjustment by cardiac response to achieve rapid replenishment of oxygen store and removal of CO_2_, as well as increases the breath events. As to active dives, meanwhile, green turtles do not deplete their oxygen stores, followed by only a few breaths for effective locomotion (Table 3, Okuyama et al., 2014). Thus, partial pressure of remaining oxygen contents in their body may not maximize tachycardia at surface after active dives.

## Acknowledgements

We gratefully acknowledge Dr. Y. Kakizoe for prescription of anesthetics and sedatives, and direction for administration of these medicines. The authors thank T. Shimizu for his kind assistance on the experiment. T. Kojima and Y. Makiguchi provided the blood gas analyzer, and useful comments on making the electrodes for ECG monitoring.

